# A calibrated functional patch clamp assay to enhance clinical variant interpretation in *KCNH2*-related long QT syndrome

**DOI:** 10.1101/2021.12.13.472492

**Authors:** Connie Jiang, Ebony Richardson, Jessica Farr, Adam P. Hill, Rizwan Ullah, Brett M. Kroncke, Steven M. Harrison, Kate L. Thomson, Jodie Ingles, Jamie I. Vandenberg, Chai-Ann Ng

**Author notes:** Co-corresponding authors Correspondence should be addressed to Jamie I. Vandenberg MBBS, Ph.D, Victor Chang Cardiac Research Institute, 405 Liverpool St, Darlinghurst, NSW 2010, Australia.; and Chai-Ann Ng, Ph.D., Victor Chang Cardiac Research Institute, 405 Liverpool St, Darlinghurst, NSW 2010, Australia.

## Abstract

**Purpose:** Modern sequencing technologies have revolutionised our detection of gene variants. In most genes, including *KCNH2*, the majority of missense variants are currently classified as variants of uncertain significance (VUS). The aim of this study is to investigate the utility of an automated patch-clamp assay for aiding clinical variant classification in the *KCNH2* gene.

**Methods:** The assay was designed according to recommendations of the ClinGen sequence variant interpretation framework. Thirty-one control variants of known clinical significance (17 pathogenic/likely pathogenic, 14 benign/likely benign) were heterozygously expressed in Flp-In HEK293 cells. Variants were analysed for effects on current density and channel gating. A panel of 44 VUS was then assessed for reclassification.

**Results:** All 17 pathogenic variant controls had reduced current density and 13/14 benign variant controls had normal current density, which enabled determination of normal and abnormal ranges for applying moderate or supporting evidence strength for variant classification. Inclusion of *KCNH2* functional assay evidence enabled us to reclassify 6 out of 44 VUS as likely pathogenic.

**Conclusion:** The high-throughput patch clamp assay can provide moderate strength evidence for clinical interpretation of clinical *KCNH2* variants and demonstrates the value proposition for developing automated patch clamp assays for other ion channel genes.

## Introduction

The revolution in genome sequencing has long promised to usher in an era of precision medicine. Whilst it is now easy to sequence and discover variants, it has proved more difficult to determine their impact on clinical phenotype. There are now more than 1 billion genetic variants that have been discovered in the human genome and are available in the build 155 version of dbSNP^1^. Of the 1.15 million variants classified in ClinVar, 41 % are listed as ‘variants of uncertain significance’ (VUS)^2^ and therefore cannot be used to aid clinical decision-making^3^. Thus there is an urgent need to develop accurate and high throughput assays to assess the functional consequences of gene variants^4^.

Under the current American College of Medical Genetics and the Association for Molecular Pathology (ACMG/AMP) variant classification framework^5^ variants are classified into one of five categories: pathogenic, likely pathogenic, VUS, likely benign or benign. All variants start as VUS and can only be actively moved towards a pathogenic or benign classification if there is sufficient evidence, based on several lines of evidence including frequency in affected individuals, segregation data, population allele frequency, *in-silico* predictions, and functional data from *in-vivo* or *in-vitro* assays. The issue of VUS is particularly acute for rare diseases, which are defined by a prevalence of less than five in 10,000^6^. By their very nature, it is rare to have clinical phenotyping data from a sufficiently large cohort of individuals with the same variant to establish strong evidence for pathogenicity. Yet, it is in rare diseases where a genetic diagnosis can be especially valuable for detecting asymptomatic cases during family screening and, in some cases, initiating preventative treatment.

Inherited long QT syndrome (LQTS) is an autosomal dominant cardiac disorder with an estimated incidence of 1 in 2000^7^. It predisposes otherwise healthy individuals to arrhythmias and sudden cardiac death^8^. A clinical diagnosis of LQTS is based on electrocardiogram (ECG) abnormalities (most noticeably a heart-rate corrected QT interval of over 470 ms in women and 450 ms in men) and suspicious personal or family history^9^. Once diagnosed, LQTS can be effectively treated with lifestyle modifications, beta blockade, cardiac denervation and, in more severe cases, with an implantable cardioverter-defibrillator (ICD)^9^. The major difficulty with managing patients and their families lies with diagnosing the disease prior to the onset of a potentially lethal arrhythmic event. This is complicated by the highly variable presentation of LQTS. For example, the hallmark QT prolongation is absent in ∼30 % of genotype-positive siblings of probands with the disease^*10*^.

The two major subtypes of LQTS are caused by variants in *KCNQ1* and *KCNH2*, which encode for cardiac potassium channels. We now have a detailed understanding of how defects in these ion channels cause LQTS^11,12^. Furthermore, ion channels are amenable to high-throughput functional analysis using automated patch clamp (APC) platforms^13–15^. Given that LQTS is (i) an actionable disease where improved diagnostic yields from genetic testing will bring tangible benefits to patients and their families; (ii) the mechanisms linking gene level defects to clinical phenotypes are well understood and (iii) there are high-throughput assays available for characterising variants in ion channels, LQTS is an excellent candidate for the development of clinically actionable functional genomics assays.

To help standardise the evaluation and application of functional evidence, the Clinical Genome (ClinGen) Sequence Variant Interpretation (SVI) Working Group recently published recommendations on validating functional assays^16^. Accordingly, the aims of this study were to validate a patch clamp assay for *KCNH2* missense variants and to determine the evidence strength that it can provide for clinical variant classification.

## Methods

### *KCNH2* patch clamp assay

#### Selection of KCNH2 variants

A total of 44 *KCNH2* (NM_000238.4) missense variants listed in the ClinVar database as either benign / likely benign or pathogenic / likely pathogenic underwent internal variant curation, by experienced cardiac genetic counsellors, using the ACMG/AMP criteria^5^. This included the following levels of evidence: absence/rarity in the population (PM2) as supporting; prevalence of the variant in affected individuals (PS4) as supporting (for 2 probands), moderate (4 probands), strong (8 probands), very strong (>16 probands); deleterious *in silico* predictions (PP3); co-segregation with disease in three or more affected family members (PP1), mutational hotspot evidence (PM1) for transmembrane/linker/pore specific for *KCNH2*^17^, appropriate allele frequencies (BA1 or BS1), multiple lines of *in silico* evidence suggested a benign effect (BP4) and multiple reputable labs had reported the variant as benign / likely benign in ClinVar (BP6). Out of the 44, 14 variants met benign / likely benign criteria and 17 met pathogenic / likely pathogenic classification without applying functional evidence (PS3)^16^. These 31 variants were used as variant controls for *KCNH2* assay validation (see Supplementary Table S1).

#### Generation of KCNH2 variant cell lines

Methods for the assay pipeline have been described in detail previously^18^. In brief, DNA plasmids containing each variant, confirmed by Sanger sequencing, were ordered from GenScript Inc (Pistcataway, NJ, USA) and subcloned into a bicistronic expression plasmid. This expression plasmid enabled co-expression of variant and wild type (WT) *KCNH2* allele to recapitulate the heterozygous expression of variants in patients with LQTS2. The plasmids were used to generate Flp-In HEK293 (Thermofisher, cat. #R78007) stable cell lines, which were assayed using an automated patch-clamp electrophysiology platform (SyncroPatch 384PE, Nanion Technologies, Munich, Germany).

#### Sample size determination for reliable current density estimation

To establish sample size requirements and the dynamic range of the assay (i.e. normal function versus complete loss of function), positive (WT Flp-In HEK293 stable cell line) and negative (Flp-In HEK293) controls were assayed across six full and two half 384-well plates, using three separate batches of cells.

#### Controls and replications

Following recommendations from the ClinGen SVI Working Group, all assay plates included WT cells as positive control and blank cells as negative controls, which left sufficient wells to assay 10 heterozygous variant cell lines per plate. Two technical replicates for each batch of cells were performed and this was repeated for a fresh batch of cells grown from a new frozen stock for the same construct. Thus each variant was assayed on 4 separate plates.

#### Data analysis

Quality control (QC) measures were applied so that only high quality patch clamp recordings were included: seal resistance > 300 MΩ, capacitance of 5-50 pF, and leak corrected current at –120 mV <40 pA of baseline^18^. A suite of voltage protocols used in patch clamp electrophysiology experiments was used to measure activation, deactivation and inactivation gating, as described in detail previously^18^. Current density was also measured by first stepping the voltage to +40 mV for 1s then measuring peak tail current at –50 mV, the voltage at which the rapid delayed rectifier current, *I*_Kr_ (which is encoded by *KCNH2*), peaks during repolarization of the cardiac action potential^11^.

The cell-cell variation in channel expression resulted in peak current densities with a non-Gaussian distribution. The peak tail current densities from each plate were transformed to a normal distribution by using a square root function, thereby allowing the use of mean and standard deviations (SD). We then normalised data to the mean value of WT from the respective plate (normalised peak tail current densities_sqrt_) to control for any plate effects. To determine the sensitivity and specificity of the peak tail current density measurements for differentiating between benign and pathogenic variants, we used 2 SD from the mean of all benign / likely benign variants to establish the threshold for functionally normal versus abnormal variants. The Odds of Pathogenicity (OddsPath) was then calculated based on the performance of the assay using the formulae derived by Brnich et al^16^, assuming a perfect binary readout.

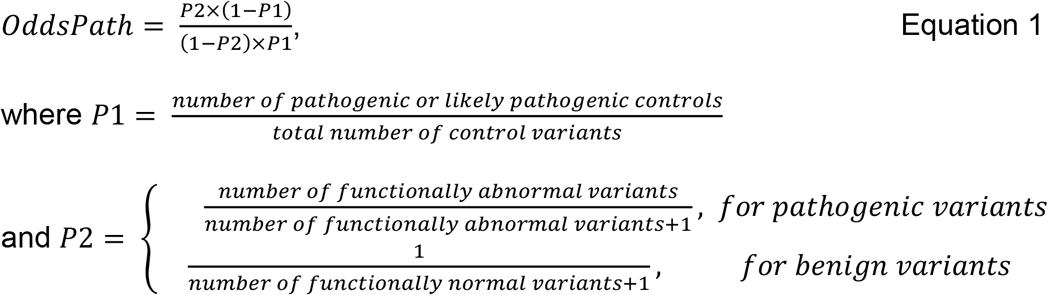

### VUS reclassification

#### Selection of variants for reclassification

A total of 45 *KCNH2* variants were selected for reclassification based on the following criteria: (1) clinical significance listed in the ClinVar database as either VUS or not provided; and (2) published functional data from a previous study^19^ indicating loss of function phenotype according to the criteria established in this study. Following baseline assessment of the 45 variants, 1 was classified as likely pathogenic and 44 were confirmed as VUS using the following criteria: absence/rarity in the population (PM2_supporting); prevalence of the variant in affected individuals (PS4) as supporting (for 2 probands), moderate (4 probands), strong (8 probands), very strong (>16 probands); deleterious *in silico* predictions (PP3); and co-segregation with disease in three or more affected family members (PP1). Functional evidence was then applied as moderate for severe loss-of-function (> 4 SD of mean) (PS3_moderate) or supporting level for partial loss-of-function (between 2-4 SD of mean) (PS3_supporting) and we assessed the resulting impact on the baseline classification.

## Results

### Sample size determination for reliable current density estimation

Figure 1Ai shows the raw current densities measured at −50 mV from 1198 recordings (from a possible 2496 wells) that passed all quality control measures (see methods). The distribution of data reflects the cell to cell variability in protein expression levels and is positively skewed (Figure 1Ai, 1Aii). The data can be converted to a normal distribution using a square root transformation. The standard deviation for this transformed distribution was 33.4 % (Figure 1Bii). This translates to requiring a minimum of 14 cells per variant to detect a 50 % difference or 54 cells per variant to detect a 25 % difference at 90 % power with 95 % confidence interval (Cl) (Supplementary Figure S1A).

**Figure 1:**
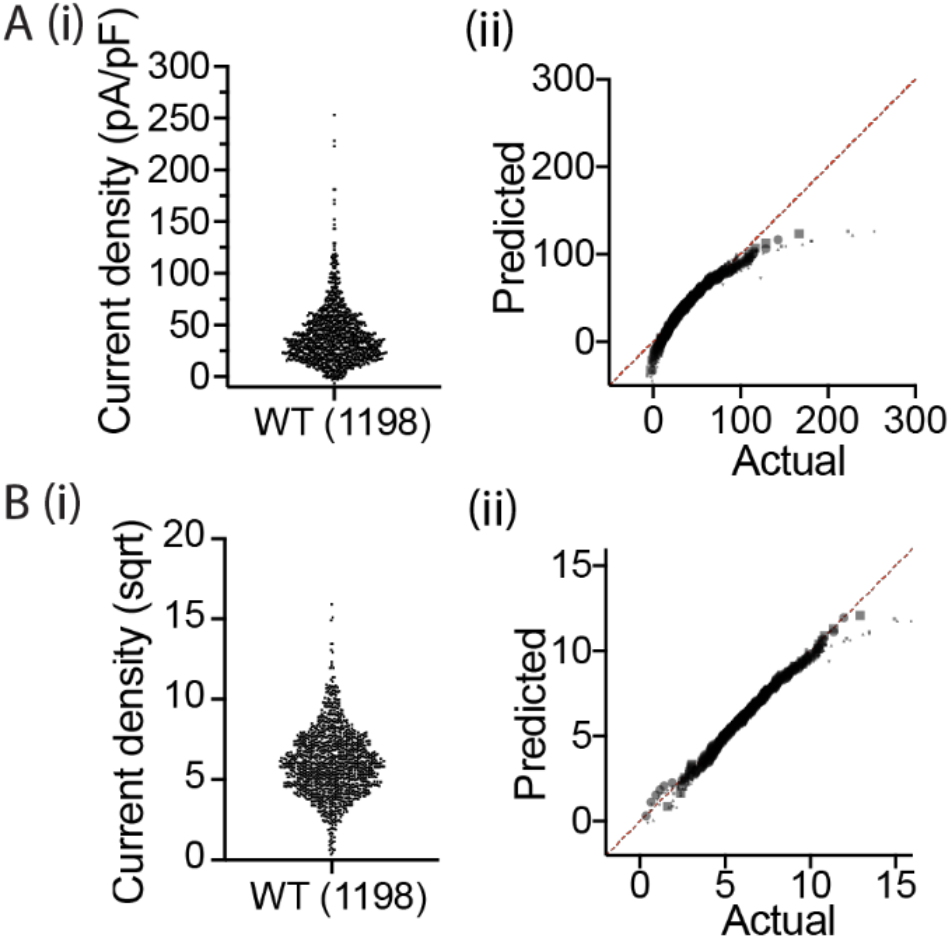
Measurement of WT current density using automated patch clamp. (Ai) Peak tail current density of WT at –50 mV for 8 different plates. (Aii) Normal probability plot showing the data is not normality distributed. (Bi) Transformation of current density in panel Ai using square root function. The standard deviation (SD) was determined to be 33.8%. (Bii) Normal probability plot after square root transformation showing the data is now normally distributed.

Based on these results we designed the assay layout to contain 32 wells for each variant, as well as 32 wells each for the WT and negative controls. Each variant was assayed on at least four different assay plates to ensure adequate sample size and reproducibility (Figure 2A). On average 67±16 (SD) recordings per variant passed all QC criteria (see Methods). Example peak tail currents for WT control wells are shown in Figure 2Bi and the peak tail current density for all WT and negative control recordings from the 24 plates are summarised as violin plots in Figure 2Bii-iii. The WT and negative controls showed a good dynamic range despite some variation between plates.

**Figure 2:**
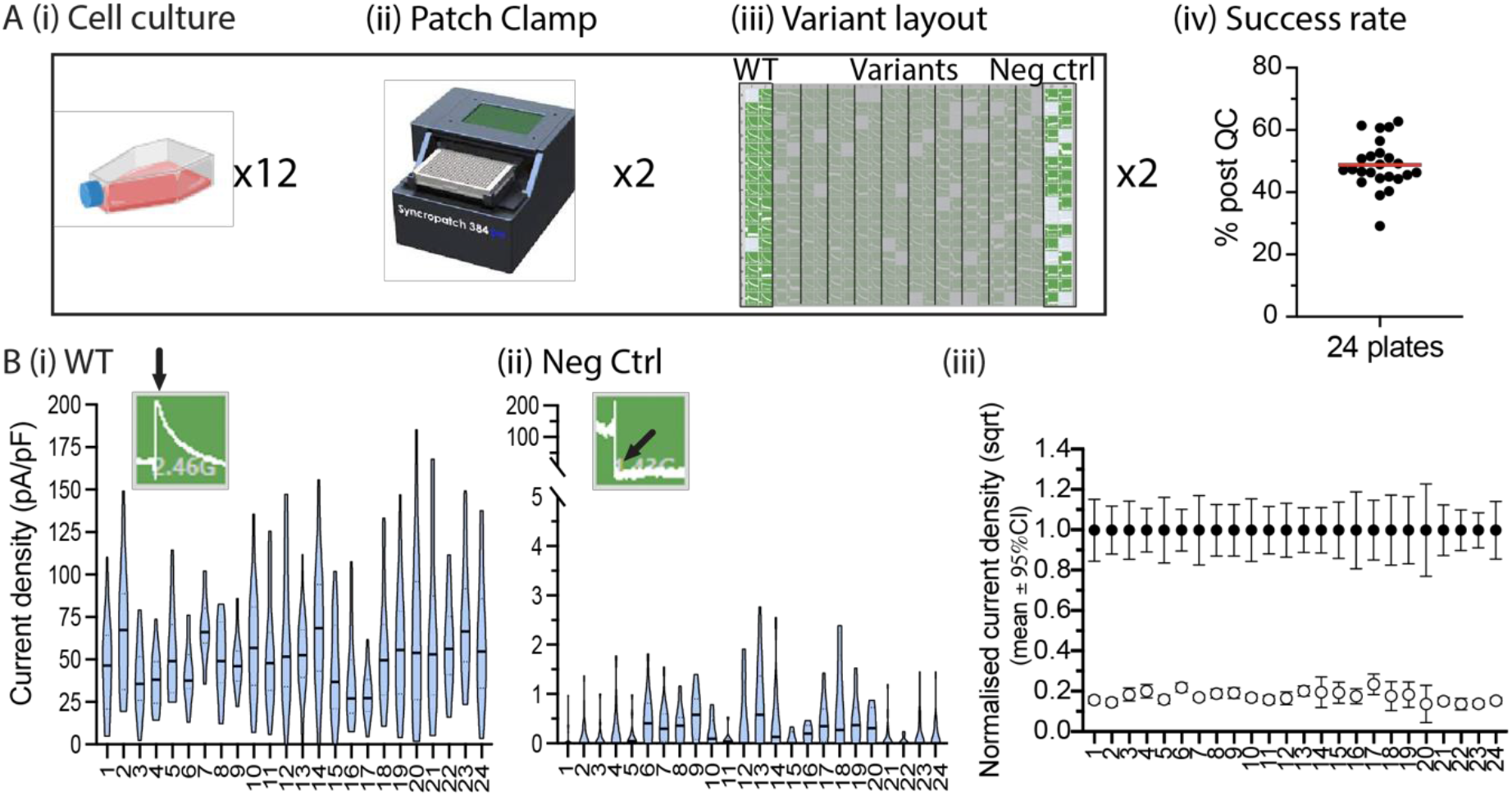
Implementation of best practice recommendations for *KCNH2* patch clamp assay. (A) Each variant was assessed 4 times (see methods) with WT and negative controls present on each plate. Overall, 49% of cells passed all quality control measures, resulting in 67 ± 16 (SD) successful recordings for each variant across the 4 plates. (Bii) Violin plot of current density at –50 mV for negative control. Insets show the peak tail current for WT and negative controls. (Biii) The dynamic range of the assay showing a clear separation between the positive (WT, filled) and negative control (clear).

### Reproducibility of the KCNH2 patch clamp assay

Example data traces and violin plot summaries for the current density measurements at −50 mV for a benign variant (p.Lys897Thr), a likely pathogenic (p.Gly584Ser) and a pathogenic (p.Ala561Val) variant are shown in Figure 3. The current density at −50 mV reflects the number of channels reaching the plasma membrane as well as impacts of variants on the activation and inactivation kinetics and any ion selectivity changes^13^. The normalised peak tail current densities_sqrt_ for these variants, presented as violin plots with each replicate plates highlighted as different colours in Figure 3B, show that the assay can detect loss of current density caused by these variants.

**Figure 3:**
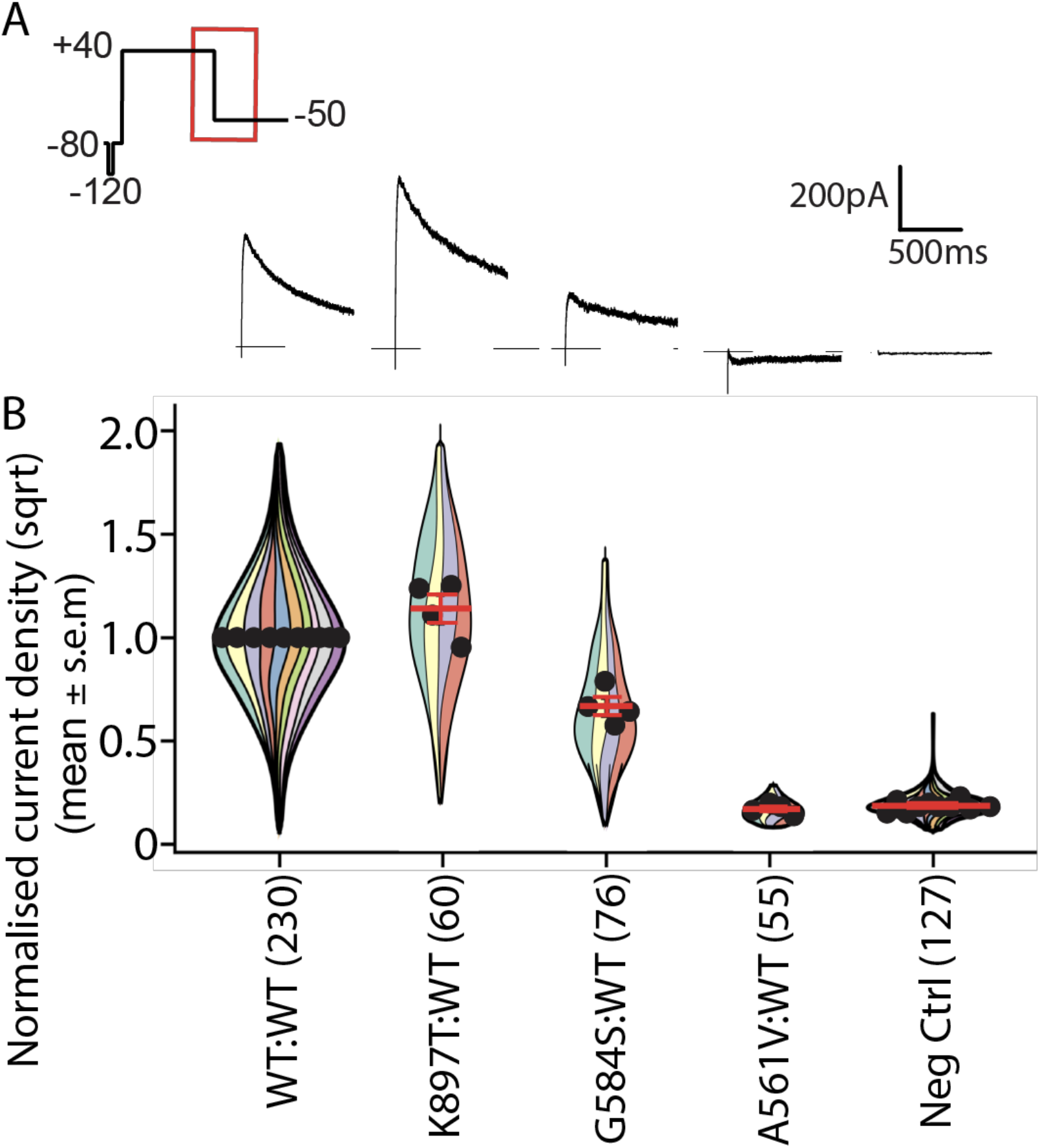
Reproducibility and replication of the *KCNH2* patch clamp assay. (A) Example peak tail current of WT:WT, p.Lys897Thr:WT (benign), p.Gly584Ser:WT (likely pathogenic), p.Ala561Val:WT (pathogenic) and negative control corresponding to the highlighted region (red box) within the voltage protocol. (B) The quantified peak tail current densities were transformed using square root function and normalised to the respective WT from the same plate. Four replicates were acquired for each variant and the corresponding WT and negative controls for those plates were also shown. The circles are mean of each replicate and error bar is the standard error of mean. The total number of patch clamp experiments from 4 replicates are indicated within the bracket.

### Determination of the functional evidence strength for KCNH2 patch clamp assay

A set of 31 variants, 14 benign / likely benign and 17 pathogenic / likely pathogenic, that had been classified in the absence of functional data (see Supplementary Table S1) was used to establish a threshold for separating functionally normal and abnormal variants. The normalised peak tail current densities_sqrt_ for the benign / likely benign and pathogenic / likely pathogenic variants were clearly segregated (Figure 4A). The functionally normal region was defined as the mean ± 2SD for the normalised peak tail current densities_sqrt_ for the WT and 14 benign / likely benign variant controls (blue region, Figure 4A). The pathogenic / likely pathogenic variant controls all reside outside the functionally normal region. Based on the threshold for normal and abnormal current density established using this set of variants, the *KCNH2* patch clamp assay achieved 100 % sensitivity and 93 % specificity (Figure 4B). The confidence interval for the mean value for p.Arg148Trp, classified as likely benign in this study using ACMG/AMP criteria, crossed over into the abnormal range. Using the formula for calculating the odds of pathogenicity provided by ClinGen SVI Working Group^16^ (see Equation 1, Methods), the assay achieved 0.063 for benign (corresponding to BS3_moderate) and 14.8 for pathogenic (corresponding to PS3_moderate; Supplementary Table S2).

**Figure 4:**
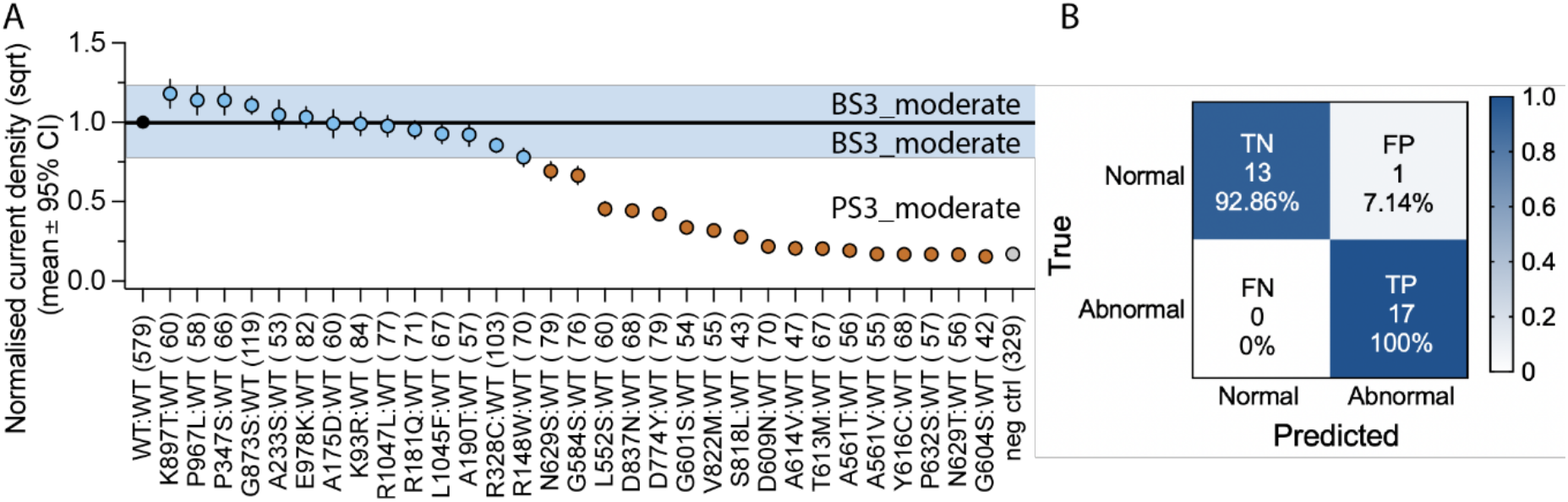
Establishment of the ACMG BS3/PS3 strength using validation variants. (A) Summary plot of the peak tail current density measurements at –50mV with square root transformation applied and normalised to WT current density on the respective plates. Data are shown as mean ± 95 % confidence interval. The mean and standard deviation of the mean for WT and the 14 benign controls (blue circles) are 1.00 ± 0.11. A variant is defined as having normal function if its mean value lies within 2 SD of the mean of all the benign/likely benign variants (highlighted by the blue region). Functionally abnormal variant for loss-of-function (brown circles) is defined as having mean value more than 2 SD below the mean of the benign variants (PS3_moderate). (B) Confusion matrix showing high specificity and high sensitivity for the classification of these variant controls based on the normal and abnormal threshold in (A).

### Application of functional data for variant reclassification

Of the 45 variants chosen for the test cohort, one was reclassified as likely pathogenic based on the current ACMG/AMP criteria while 44 remained as VUS before applying functional data (Figure 5A; Supplementary Table S3; column P). Analysis of current density data for these 44 variants supported application of PS3_moderate for 31/44 VUS, and PS3_supporting for 13/44. Integrating this functional data with existing evidence, allowed 6/44 VUS to be reclassified as likely pathogenic (Figure 5B; Supplementary Table S3; columns U/V). If the point-based system for variant classification proposed by Tavtigian and colleagues^20^ were to be adopted, then an additional 9 VUS could be reclassified to likely pathogenic (Figure 5C).

**Figure 5:**
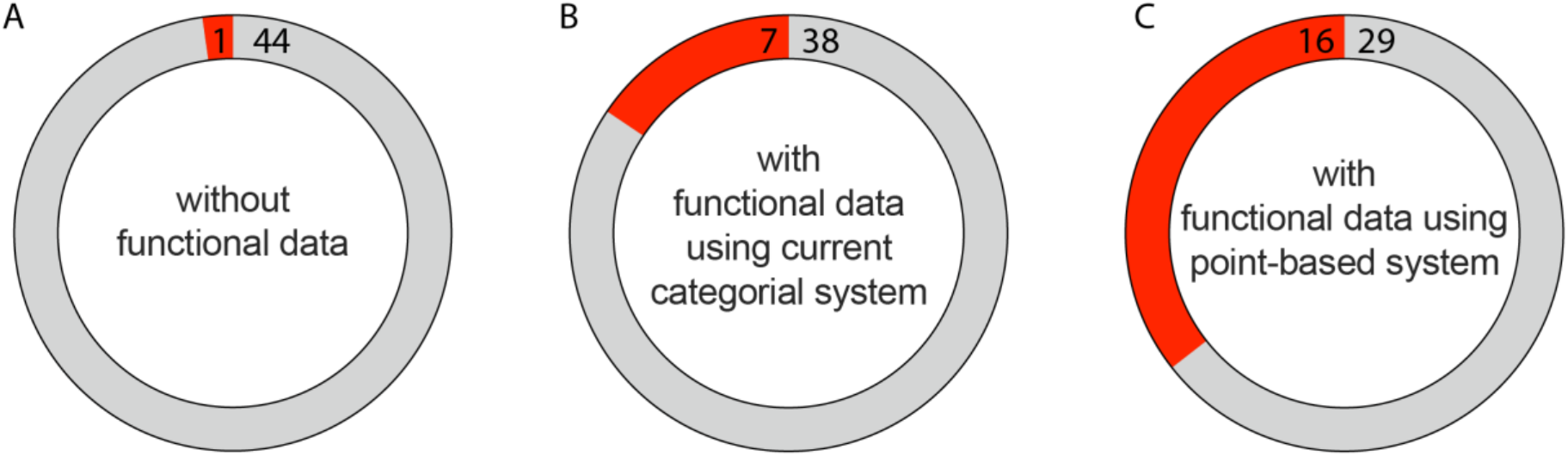
Reclassification of *KCNH2* VUS that have abnormal function. (A) Categorial classification without functional data. (B) Categorial classification with functional data applied as moderate or supporting (Supplementary Table S3). (C) Point-based classification^20^ with PS3_moderate being applied as 3 points. Red is likely pathogenic and grey is VUS.

## Discussion

Patch clamp assays have been a mainstay for assessing the functional impact of genetic variants in ion channel genes, including *KCNH2* (reviewed in Delisle et al^21^). Traditionally, these assays have been very labour intensive with typical throughputs of 10-20 cells per day. However, with the development of automated patch clamp platforms this throughput has increased to hundreds-thousands of cells per day^22^, thus making it feasible to start addressing the deluge of VUS in ion channel genes. Here, we have formally validated a *KCNH2* automated patch clamp assay based on the recommendations proposed by the ClinGen SVI Working Group^16^. Following recommendations and results of our preliminary study we designed the assay to incorporate the following: (1) inclusion of positive and negative controls on each 384-well plate; (2) allocation of 32 wells per variant on at least four different assay plates to ensure adequate sample size and reproducibility; (3) normalisation to WT (positive control) on each plate and (4) transformation of data to ensure a normal distribution of the data. Using a cut off between functionally normal and functionally abnormal as two SD below the mean value of the 14 benign / likely benign variant controls, the assay produced concordant results for 13/14 benign / likely benign variants and 17/17 pathogenic / likely pathogenic variant controls. Based on the Odds of Pathogenicity (OddsPath) statistical analysis developed by Brnich et al., our assay is currently capable of providing functional evidence at moderate level. However, for those variants that were only 2-4 SD lower than the mean of the benign variants we have adopted a more conservative classification of supporting level evidence as recommended by Brnich et al for variants with partial loss of function^16^. With this evidence we could reclassify 6 out of 44 VUS (13.6 %) to likely pathogenic.

### Current density at –50mV captures the phenotype of KCNH2 variants

Variant function was determined by measuring peak tail current density at –50mV, a measure which reflects (i) the number of channels that have trafficked to the plasma membrane, (ii) the proportion of channels that are activated during the depolarised voltage at +40 mV, (iii) the proportion of channels that have recovered from inactivation at –50 mV^18^ and (iv) the ionic selectivity and hence reversal potential for current flow. The vast majority (>80%) of loss-of-function missense variants in *KCNH2* that have been characterised to date are known to reduce current density via trafficking defects. There are, however, a small number of variants that reduce activation^23^, enhance inactivation^24^ or alter ion selectivity^13^ which will also be detected as reduced current density in our assay. Thus, this single assay interrogates the vast majority of mechanisms by which missense variants can affect HERG current levels^21^.

For completeness, we further investigated channel gating in variants with sufficient current density. We did not detect abnormal gating in any of the benign / likely benign variant controls (Supplementary Figure S2). The assay, however, was able to detect known gating defects in likely pathogenic variant controls, i.e. the inactivation gating defect in p.Gly584Ser^24^ and deactivation defect in p.Leu552Ser^25^. It is also notable that the current density values across the cells were not normally distributed, which is consistent with what has been seen for cell-cell variation in single cell gene expression studies^26^. The positively skewed raw data could be converted to a normal distribution by applying a square root transformation and so this transformation was applied to all current density measurements.

Based on our analysis of the 14 benign / likely benign variant controls, the lowest threshold to be considered as functionally normal was 0.78 (for square root of current density), which is equivalent to 61 % of WT current density (pA/pF). This is slightly higher than the predicted 50 % loss-of-function caused by heterozygous nonsense variants that are well established for causing LQTS2, albeit with incomplete penetrance^10^.

### Functional patch clamp assay has high sensitivity and specificity

All pathogenic / likely pathogenic variant controls fell in the functionally abnormal range. This indicates that our *KCNH2* patch-clamp assay is highly sensitive for identifying functionally abnormal variants. It is also highly specific, which is useful for discerning the status of variants that are identified as incidental or secondary genetic findings. However, there is a caveat that assays based on transfected cDNAs in heterologous expression systems may miss variants that affect splicing or interactions with other proteins. The one likely benign variant that had its confidence interval fall outside the normal range was p.Arg148Trp. This variant was classified internally as likely benign mainly due to its high minor allele frequency (gnomAD V2.1.1 AC: 0.001133; gnomAD V3 AC: 0.0006441). However, there are conflicting classifications in ClinVar (Benign(5); Likely benign(3); Uncertain significance(2)) and there are numerous reports linking p.Arg148Trp to LQTS2^27–29^. Thus it is possible that p.Arg148Trp, while insufficient to cause LQTS2 alone, does have a deleterious effect on function, which maybe a risk allele^30^.

### Incorporation of functional data for variant classification in long QT syndrome genes

Automated patch clamp data can provide functional evidence for *KCNQ1, KCNH2* and *SCN5A* variant classification^13–15^. However, factors such as a lack of detailed guidance on how functional evidence should be evaluated, and differences in the application of the functional evidence strength (PS3/BS3) criterion has contributed to discordances in variant interpretation between diagnostic laboratories^16^. The *KCNH2* patch clamp assay reported in this study was stringently assessed to provide functional evidence for normal and abnormal at moderate levels (BS3_moderate or PS3_moderate). However, given the inherent limitations of *in vitro* functional assays and the potential lethality of LQTS2, it may be appropriate to take a more conservative approach. For example, having normal *in-vitro* function does not exclude the possibility that a variant might be abnormal *in-vivo, e*.*g*. HEK cell assay may not fully recapitulate all the protein interactions that occur in native cardiomyocytes. We accordingly suggest that the evidence provided for functionally normal variants should be applied at supporting level (BS3_supporting) when classifying clinical *KCNH2* variants. For functionally abnormal variants, we suggest applying evidence at moderate level (PS3_moderate) for *KCNH2* variants with severe loss-of-function (>4 SD from the mean of benign / likely benign variants) or supporting level for *KCNH2* variants with partial loss-of-function (2-4 SD from the mean of benign / likely benign variant controls) (Supplementary Table S3; column R).

One limitation/barrier to developing functional assays for assessing LQTS variants is the lack of definitively benign variants in LQTS genes (*KCNH2*, as well as in *SCN5A* or *KCNQ1)*, which restricts the strength of evidence that can be provided by a functional assay to a moderate level (PS3_moderate)^31^. Previous studies have used patch clamp data at strong level evidence for variant classification^15,32,33^ although this contradicts the advice in current ACMG/AMP categorial classification system. The OddsPath ratio for our *KCNH2* assay for PS3 is 14.3:1, which does not quite reach the level required for a strong level of evidence (minimum OddsPath of 18.7:1 for strong and 4.3:1 for moderate). If a point-based classification system^20^ was used for PS3, our assay would be given three points rather than 2 (moderate) or 4 (strong) which would result in an additional 9 VUS being reclassified as likely pathogenic. The evaluation of a larger number of benign variants, which could occur once a larger proportion of the population have had their genomes sequenced and more benign variants are identified, will very likely allow the evidence strength of the assay to increase to a strong level (PS3/BS3) and thus increase the clinical utility of the assay.

### Significance

Deciphering the clinical impact of VUS is a key step to unlocking the full potential of genomic medicine. High-throughput functional assays, such as the *KCNH2* patch clamp assay described here, can improve the diagnostic yield of genetic testing. Our result has important implications for the management of patients with LQTS2 in terms of diagnosis and the provision of gene-specific advice on lifestyle modifications and family screening^9,34^. Results from this assay will be an important resource for the ClinGen Variant Curation Expert Panel (VCEP) when establishing gene-specific criteria for *KCNH2*. Our work also provides a blueprint for the validation of future automated patch clamp assays in other ion channel genes.

## Acknowledgement

This project was funded by the Australian Genomics Cardiovascular Genetic Disorders Flagship (funded through the Medical Research Future Fund to JIV, CAN and APH), a NSW Cardiovascular Disease Senior Scientist Grant (JIV), an National Health and Medical Research Council Principal Research Fellowship (JIV), National Institutes of Health grant R00HL135442 (BMK), Leducq Foundation for Cardiovascular Research grant 18CVD05 “Towards Precision Medicine with Human iPSCs for Cardiac Channelopathies” (BMK), American Heart Association Career Development Award 848898 (BMK). National Health and Medical Research Council Career Development Fellowship #1162929 (JI). We also acknowledge support of the Victor Chang Cardiac Research Institute Innovation Centre, funded by the NSW Government.

## Supplementary Figures

**Supplementary Figure S1:**
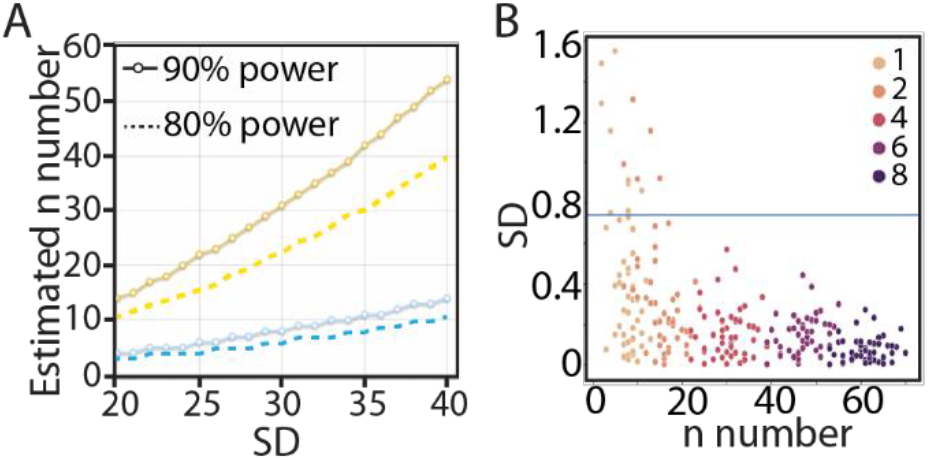
Power analysis and sample size determination. (A) Estimated n number required for detecting 25 % or 50 % difference at different standard deviations. Dashed and solid lines are the n number required for achieving 80 % and 90 % power with 95 % confidence interval, respectively. (B) The mean of the subgroup was then compared with the population mean to calculate the difference relative to the population mean. This was repeated 50 times for subgroups containing 1, 2, 4, 6 and 8 columns of data.

**Supplementary Figure S2:**
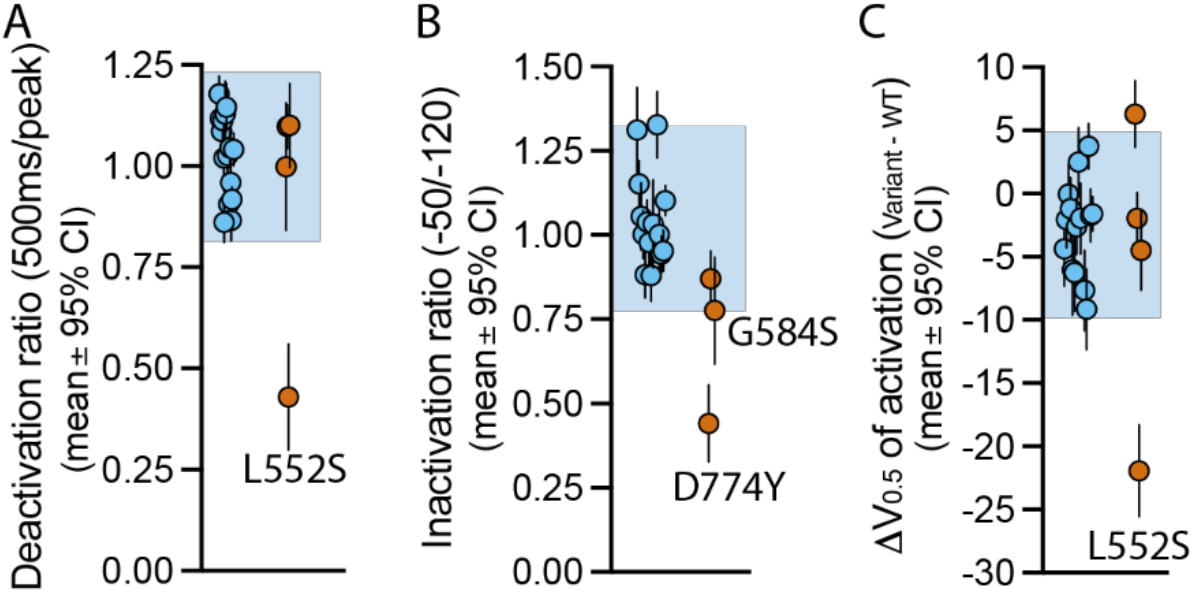
Establishment of functional normal range for channel gating. (A) Deactivation ratio of current amplitude (500ms after peak tail current / peak tail current) measured at –50 mV. (B) Steady-state inactivation measured as the ratio between the peak tail current amplitude at –50 and –120 mV. (C) The *V*_0.5_ of activation presented as the difference from WT. The functionally normal range was established to be within the 2 SD (blue region) of the mean for the values obtained from the 15 benign / likely benign variant controls (blue circles). Only pathogenic / likely pathogenic variant controls that have sufficient current density were analysed for gating phenotype (brown circles).

